# decOM: Similarity-based microbial source tracking of ancient oral samples using k-mer-based methods

**DOI:** 10.1101/2023.01.26.525439

**Authors:** Camila Duitama González, Riccardo Vicedomini, Téo Lemane, Nicolas Rascovan, Hugues Richard, Rayan Chikhi

## Abstract

**Background:** The analysis of ancient oral metagenomes from archaeological human and animal samples is largely confounded by contaminant DNA sequences from modern and environmental sources. Existing methods for Microbial Source Tracking (MST) estimate the proportions of environmental sources, but do not perform well on ancient metagenomes. We developed a novel method called decOM for Microbial Source Tracking and classification of ancient and modern metagenomic samples using k-mer matrices.

**Results:** We analysed a collection of 360 ancient oral, modern oral, sediment/soil and skin metagenomes, using stratified five-fold cross-validation. decOM estimates the contributions of these source environments in ancient oral metagenomic samples with high accuracy, outperforming two state-of-the-art methods for source tracking, FEAST and mSourceTracker.

**Conclusions:** decOM is a high-accuracy microbial source tracking method, suitable for ancient oral metagenomic data sets. The decOM method is generic and could also be adapted for MST of other ancient and modern types of metagenomes. We anticipate that decOM will be a valuable tool for MST of ancient metagenomic studies.

## Background

Ancient metagenomics is the study of multi-species genomic data from samples that have degraded over relatively long time [1]. Analysing ancient DNA (aDNA) is particularly challenging due to deterioration and contamination with environmental and modern contaminant DNA sequences. Deterioration refers to DNA damage, which in genetic material from fossil records usually comes in the form of depurination, nick formation and cytosine deamination [2]. Contamination refers to genetic material (ancient or modern) that does not derive from the sample of interest [3]. It can come from the microbes that are present in decaying tissue, from the soil or sediment where the samples were taken, or be an unintended consequence of manipulation during and after excavation [4, 5]. Despite following well-established standards and precautions to prevent modern DNA contamination and reduce the proportion of environmental microbial taxa [5, 6], a certain level of unwanted genetic material in the samples is unavoidable [4]. Under these circumstances, contamination assessment of aDNA samples is crucial not only to avoid misleading results after downstream analysis, but also to decide which samples are worth to be further sequenced [7].

The task of Microbial Source Tracking (MST) is to quantify the proportion of different microbial environments (sources) in a target microbial community (sink) [8]. MST allows to quantify contamination [9] in metagenomics sequencing data and to predict the metadata class of a given microbial sample.

Two of the most widely used methods today for MST in metagenomic data are metagenomic-SourceTracker (mSourceTracker)[10] and FEAST [8], which depend on previously annotated data using taxonomic abundance profiles. mSourceTracker is a metagenomic extension of the popular SourceTracker [9], a method that estimates contamination proportions using a mixture model of taxonomic profiles via Gibbs sampling. It is known that the sensitivity of SourceTracker can be improved through parameter adjustments [11], however more rigorous evaluations are still needed to fully understand the effect of adjusting multiple parameters and hyperparameters on its performance [12]. FEAST, released 8 years after SourceTracker, uses an expectation-maximisation approach that reduced the running time of SourceTracker by a factor of 30 or more. It has been reported to require parameter tuning to achieve optimal performance [13], which is a resource intensive procedure when handling with large data sets.

FEAST and mSourceTracker require a reference database which is necessary to build the taxonomy-based clustering tables that both methods use as input. Indeed, in both cases, metagenomic data must be grouped into bins or clusters of sequences sharing the same taxonomic classification, an information that is not only highly dependent of the database used, but also highly biased by the limited proportion of the microbial diversity that has been already sequenced and taxonomically annotated. [14].

Finally, these taxonomy-based clustering tables can also lead to misleading results depending on the sequence similarity metric and the threshold used to define them [15]. To our knowledge, there are no reported reference-free methods for contamination assessment that use MST for large-scale metagenomic analyses [13]. In this work we seek to move away from database-dependent methods and use unsupervised approaches exploiting read-level sequence composition and the wealth of information contained in metagenomes that were previously sequenced.

Over the past years and with the decrease of sequencing costs, large databases of metagenomic collections from all sorts of environments have become available [16, 17, 18]. These metagenomic raw reads collectively require petabases of storage, which prohibits their reanalysis by most labs. This prompted the development of efficient methods for exploring the sequence information contained in these collections, via searching substrings of length k (k-mers) [19]. Such methods build an index of all k-mers and their counts over a collection of samples in the form of a k-mer matrix, where each cell of the matrix represents the abundance (or presence/absence) of a k-mer in a sample. Such matrices are a concise representation of genomic data that deals more efficiently with sequencing errors and genetic variation [19]. Tools such as kmtricks [20] allow the rapid construction of k-mer matrices from massive collections of sequencing data sets.

In this study we developed a novel reference-free and k-mer-based method called decOM to perform MST and environmental type prediction of a given microbial sample. decOM was evaluated in a collection of ancient oral metagenomes with variable contamination levels. Our results show that decOM outperforms two of the most commonly used MST methods in the multi-class classification task of finding the most abundant source environment in a sink. We tested our methodology on a collection of 360 metagenomic data sets of ancient oral samples and its possible contaminants, in an external validation set of 254 ancient oral samples and on a simulated ancient calculus metagenome.

## Implementation

### Evaluation setting

Dental calculus or tartar is mineralized dental plaque that contains remnants of microorganisms located in the oral cavity [3], and has been established over the past few years as one of the richest sources of aDNA in the archaeological record [21]. Ancient dental calculus is a great source of biomolecules (including genetic material) that originate from the host, microbes, food and the environment [6]. Dental calculus is an important reservoir of ancient human oral microbiomes, and it offers a unique possibility to examine the links between human health, diet, lifestyle and the environment throughout the course of human evolution [22]. Due to the proven relevance of aOral samples isolated from calculus in the field of ancient paleogenomics, we decided to perform our evaluations on a collection of aOral metagenomic samples and their possible sources of contamination.

The microbial composition of a given aOral sample isolated from dental calculus has been modelled in previous studies as a mixture of DNA originating from dental plaque, skin bacteria, soil and other sources [23, 24]. For this reason, we gathered 360 metagenomic data sets of diverse environment types: ancient oral (aOral), sediment/soil, skin, or modern oral (mOral) (Figure 1). We used this collection of real metagenomic data to model the contribution of possible contaminants coming from sediment/soil and skin sources in a group of aOral samples. In addition, we included a set of mOral samples to assess whether our method can tell apart modern and ancient oral environments.

**Figure 1:**
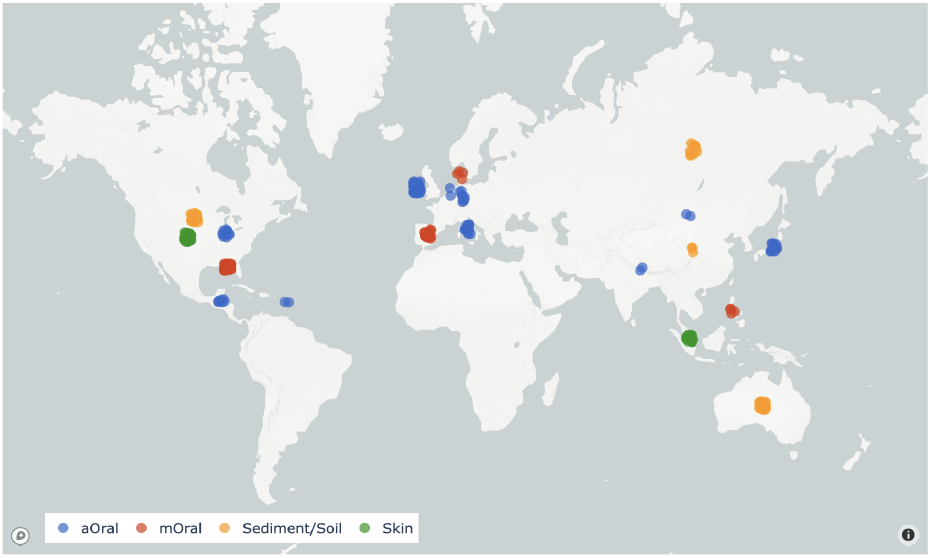
Geographical location of samples coloured by environmental type. Labels for each sample were retrieved from their metadata. The final collection of metagenomic samples included 116 (32.2%) aOral, 81 skin (22.5%), 79 sedi-ment or soil (21.9%) and 84 mOral (23.3%) samples.

The run accession codes for every aOral sample were retrieved from AncientMetagenomeDir [1], a community-curated collection of annotated ancient metagenomic sample lists and standardised metadata. Samples other than aOral were selected either because they had been used by competing MST methods or because they were labelled as aforementioned classes in well-known metagenomic databases such as curatedMetagenomicData [25], the HumanMetagenomeDB [26] or MGnify [27].

We rely on the metadata of each metagenomic sample to assign a true label (i.e. environment type), however, there is no ground truth as to what is the true proportion of aOral, mOral, sediment/soil or skin content in any of them. Several variables accessible through the metadata of each run accession are plotted in the Supplementary File (Figures 1, 2, 3 and 4).

**Figure 2:**
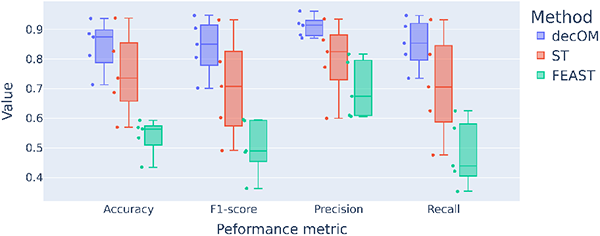
Bioproject stratified 5-fold cross-validation performance of every method. The performance from every fold was evaluated using accuracy, precision, recall and F1-score

**Figure 3:**
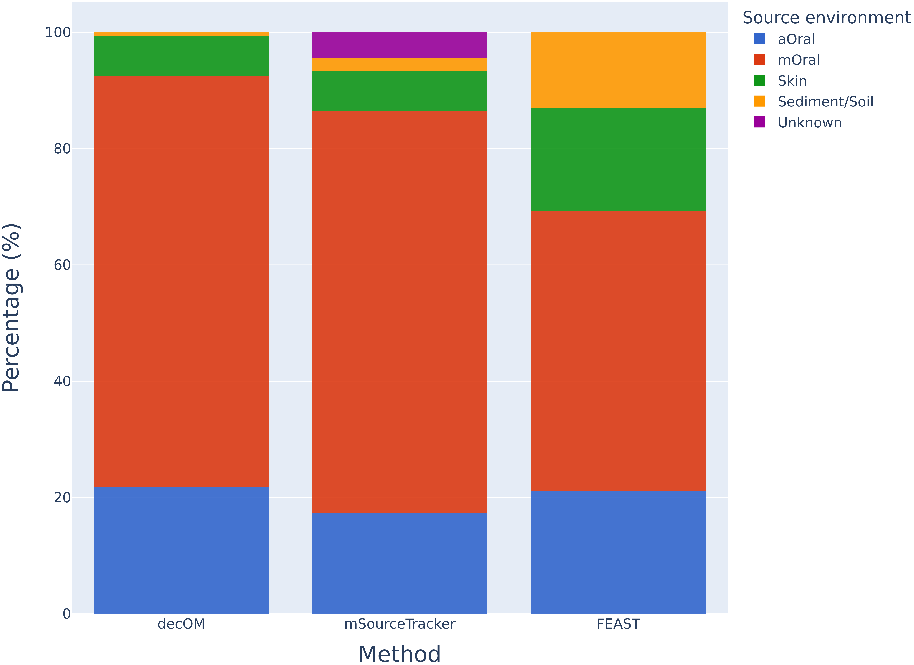
MST on a simulated ancient dental calculus metagenome. Bar plots for the source environment proportion estimation obtained after evaluating each method using as sources all the samples from the 360 metagenomic collection, and using as sink a synthetic ancient oral data set. The expected content of this synthetic sample is 100% oral

**Figure 4:**
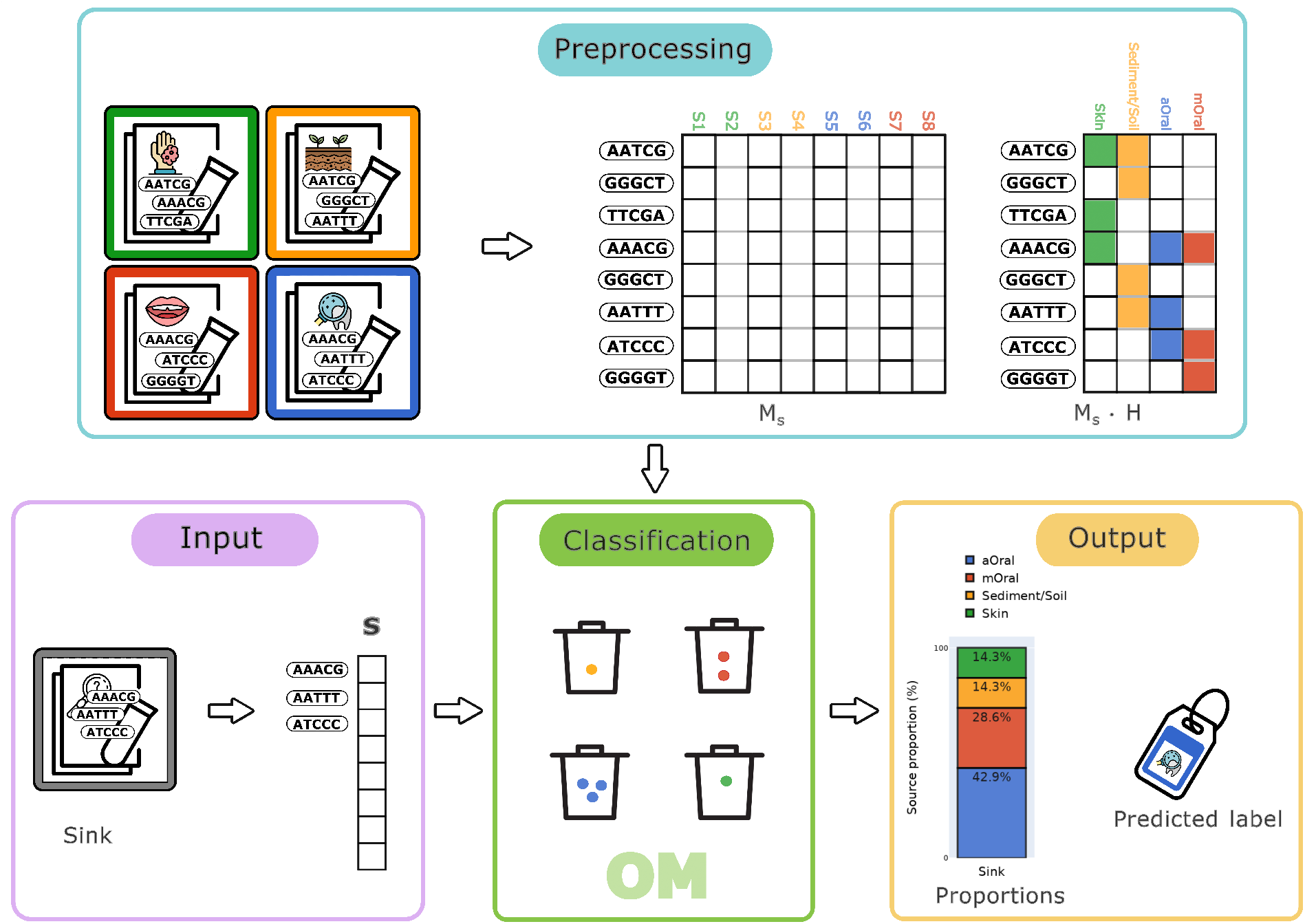
Graphical representation of decOM. Our method preprocesses an input k-mer matrix of aOral metagenomic samples and its possible contaminants, divides it into sinks and sources and then estimates and outputs the proportions of each source environment in the sink. The core idea in the classification step is that if a k-mer is present in the sink *s* represented by the vector **m**^(*s*)^, and in the source vector **m**^(*j*)^ with environment label *l*_*j*_, then a ball is added to the bin with label *l*_*j*_ (Ex: K-mer AAACG is present in the input sink S and in source S1 labelled as skin, S5 labelled as aOral and S7 labelled as mOral, hence one ball is added to the bin of skin, aOral and mOral respectively). After every entry in the the sink vector is compared against every entry of every vector in the sources, decOM outputs the estimated environment proportions and the hard label assigned to the sink *s* is that of the environment with the highest contribution.

### Input data

Both mSourceTracker and FEAST require taxonomy-based clustering tables as input. We built these tables using Kaiju [28] and the reference database NCBI BLAST nr+euk (2021-02-24 release), a non-redundant protein database of bacteria, archaea, viruses, fungi, and microbial eukaryotes (information to download it in Supplementary File, Section 1)

On the other hand, decOM takes as input a binary k-mer matrix of distinct k-mers across a collection of metagenomic samples. We used kmtricks (v1.1.1) to build a presence/absence k-mer matrix from the 360 metagenomic samples in the collection. In order to find patterns that helped us distinguish between samples from different source environments, we kept only k-mers that were present in at least 3 samples in the collection. The k-mer size in kmtricks was set to 31. We removed all k-mers seen only once in a sample, which were likely to be sequencing errors. The rest of the parameters of kmtricks were set by default.

The complete k-mer matrix contains around 9 billion k-mers, represented by 700 disjoint sets of k-mers called *partitions*. Omitting some technical aspects [29] for clarity, partitions can be seen as a random subset of the rows of the k-mer matrix, created to avoid loading the entire matrix in memory [20]. In this work we configure kmtricks to only construct a single partition out of the 700, i.e. we consider only a subset of around 14 million k-mers (0.1% of total) for subsequent analysis. We also tested with 7 partitions (Figures 13 and 14 in Supplementary File), and while it improves results marginally, the marked performance improvement when using only 1 partition justifies keeping this regime.

### Mathematical formulation

We consider a binary k-mer matrix *M* (as output by kmtricks) that indicates the presence/absence of each k-mer found across several metagenomic data sets, with *N* number of unique samples (columns) and *K* number of unique k-mers (rows). Each sample *j* is represented by a column vector **m**^(*j*)^ = (*m*_1*j*_, *m*_2*j*_, *m*_3*j*_, …, *m*_*Kj*_) where *m*_*i,j*_ corresponds to the presence/absence of k-mer *i* in sample *j*. We will use the terminology of *sink* and *sources* to respectively denote the sample we want to evaluate the composition of, and the set of samples used as a database.

Consider that the matrix *M* contains jointly all sources and potential sinks. Let a sample *s* (where *s* ∈{ 1, 2, …, *N*}) be a sink and **m**^(*s*)^ be its column vector. A source is a collection of *L >* 0 column vectors used to build a matrix of sources *M*_*s*_ of dimensions *K* × (*L* − 1). Each column vector in the sources matrix *M*_*s*_ has an associated label that comes from a finite ordered set of environments (classes) *C* = {*c*_1_, *c*_2_, *c*_3_, …, *c*_*n*_ } determined by the user. In our case |*C*| = 4, as *C* = {*aOral, mOral, skin, sediment/soil*}. The vector of labels for each sample in the sources of length *L* − 1 is represented by *ℓ* = (*ℓ*_1_, *ℓ*_2_, *ℓ*_3_, …, *ℓ*_*L*−1_), and each entry of the vector can only take one of the values from *C* as in a multi-class classification problem. The vector of categorical labels *ℓ* can be further encoded as a highly sparse one-hot binary matrix *H* of size (*L* − 1) × |*C*| where :

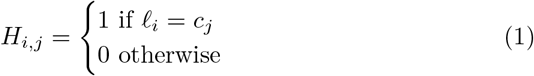

Making an analogy with bins (source environments) and balls (k-mers present in a certain source environment), we are interested in counting the number of balls that fall into each bin. The core idea of decOM is that if a k-mer is present in the sink represented by the vector **m**^(*s*)^ *and* in the source vector **m**^(*j*)^ with environment label *ℓ*_*j*_, then a ball is added to the bin with label *ℓ*_*j*_. We then compare the sink vector **m**^(*s*)^ against every source vector until all sources are exhausted. The output of this comparison is the vector **w** of length | *C*|, where every entry corresponds to the total number of balls in a certain bin, that is, the contribution of each source environment to the sink *s*.

Counting k-mers of sinks in sources amounts to performing the following matrix vector operation:

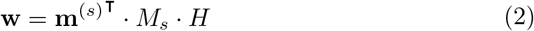

In order to produce proportions instead of raw counts, we estimate the percentage based on the total number of balls counted per bin (of all known sources)

Such proportions correspond to every element in the vector **p** = ⟨*w*_1_, *w*_2_, *w*_3_…, *w*_|*C*|_ ⟩ when multiplied by a scalar, as seen in the following operation:

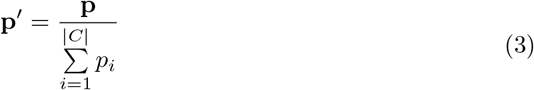

To analyse a new metagenomic sample, it is only needed to compute a presence/absence vector of k-mers for this sample using kmtricks, then this new sink is compared against the pre-computed collection of sources. decOM incorporates a kmtricks module so that the user can give as input a simple FASTQ/FASTA file of their sink of interest, rather than a presence/absence vector. Figure 4 provides a graphical representation of our pipeline.

Finally, we are working to include the contribution of an unknown source by characterising it as the number of k-mers that are present in the sink and absent in ***all*** of the sources.

decOM was implemented in Python 3.6 as a conda package and the installation instructions are available in a GitHub repository[30].

### Microbial Source Tracking evaluated in four different experimental settings

We perform a metagenomic Microbial Source Tracking to benchmark decOM, mSourceTracker, and FEAST, which all rely on an input matrix. For mSourceTracker and FEAST the input matrix corresponds to a taxonomy-based clustering table, whereas decOM takes as input a binary k-mer matrix across metagenomic data sets.

Consider the set *X* = {**m**^(1)^, **m**^(2)^, **m**^(3)^, …**m**^(*N*)^ }, where *X* contains all the column vectors of the aforementioned k-mer matrix. Let *A* = { **m**^(*s*)^ } be a set of sink vectors, and *B* = {*X*\ **m**^(*s*)^ }a set of sources. In order to estimate the proportion of source environments in each data set in our collection we run our method in a leave-one-out fashion, i.e., every run of our method uses one different sample as sink and leaves the rest of the samples as sources. One run of this experimental setup is described by Algorithm 1. Additionally, we performed a 5-fold cross-validation experiment by splitting the collection of metagenomic samples into 5 stratified folds with non-overlapping groups. The groups were defined by the BioProject from which each data set originated. A BioProject is a collection of biological data related to a single initiative originating from a single organisation or from a consortium [31]. The folds were made trying to preserve the percentage of samples for each class, given the constraint that the same group (BioProject) will not appear in two different folds. The idea behind this additional group stratification is to account for the possible bias that might appear when classifying a sink that is very similar to a set of sources simply because they come from the same BioProject and not because there is an underlying sequence similarity between the samples.

#### Algorithm 1

Pseudocode of our method used to estimate proportions of sources in sink *s*

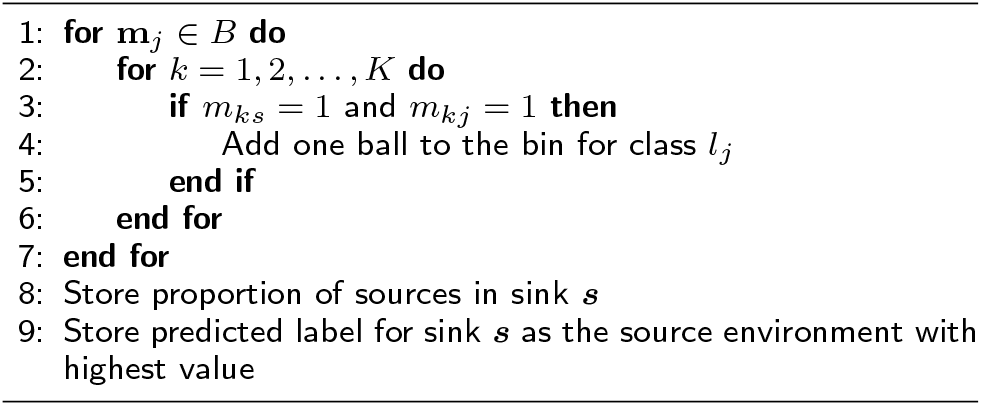

For the leave-one-out and cross-validation experiments we evaluated all methods using the Reciever Operating Characteristic (ROC) and Precision-Recall curves, and a hard label was set using as threshold the environment class with the highest contribution to the sink. Performance metrics used were Accuracy, Precision, Recall and F1-score as they are implemented in scikit-learn [32]. Because the framework of evaluation was a multi-class classification task, the performance metrics reported here were estimated for each label and then averaged across classes. Definitions for each performance metric used are specified in Section 5 of the Supplementary File.

We also tested decOM on a validation set of 254 aOral samples, none of which belonged to the collection of 360 samples we used to construct the k-mer matrix. For this experiment, the aforementioned matrix is used as sources, whereas the 254 external aOral samples are used as sinks. Because all samples belong to the same class, Precision and F1-score are not well-defined, whereas Recall and Accuracy are equivalent (See Section 5 in Supplementary File), which is why performance is measured using Recall only. Finally we tested decOM and its competitors on an uncontaminated simulated ancient oral data set and presented the estimated proportions.

## Results

We created decOM as reference-free and open-source Microbial Source Tracking method that is adapted to ancient metagenomic experiments. Our method takes as input a set of source vectors in the form of a presence/absence k-mer matrix (built from a collection of metagenomic data sets ready for the user to download), and one or more FASTA/FASTQ files to be used as sinks. It outputs a set of proportions (percentages) and a predicted metadata class per sink.

### decOM robustly predicts metagenome sample labels

#### Leave-one-out experiment

We compared the performance of decOM with FEAST [8] and mSourceTracker [9] based on their ability to correctly predict the environmental type of a sample, defined as the highest proportion among the four possible sample types (ancient oral, model oral, skin, soil). For all methods, we used the same collection of 360 metagenomic experiments as sources.

All methods output a set of proportions for each sample. We ran them in a leave-one-out fashion (one sample was used as sink, and the rest were left out as sources). In order to perform a multi-class classification task, we mapped the set of continuous proportions into a hard label, by simply assigning a label to the sample corresponding to the environmental type with the largest proportion among all the predicted sources. The performance metrics presented were calculated using the hard labels.

Table 1 shows that decOM outperforms both mSourceTracker (+3% Accuracy, +8% Precision, +3% Recall, +5% F1 score) and FEAST (+19% Accuracy, +37% Precision, +12% Recall, +33% F1 score) in the multiclass classification task of predicting source environment with the largest contribution in a sink, when such contribution is estimated using a MST framework. Precision-Recall and ROC curves are shown in the Supplementary File (See Figure 10 and 11).

**Table 1:** Environment type prediction performance of decOM, FEAST and mSourceTracker. Accuracy, precision, recall and F1-score for were estimated as an average across all classes in a leave-one-out fashion.

| Method | Accuracy | Precision | Recall | F1-score |
| --- | --- | --- | --- | --- |
| decOM | **0.8703** | **0.9184** | **0.8703** | **0.8753** |
| FEAST | 0.6816 | 0.5516 | 0.7452 | 0.5479 |
| mSourceTracker | 0.8388 | 0.8388 | 0.8388 | 0.8289 |

#### Cross-validation

To further validate that decOM does not solely rely on closely related samples for its predictions, we performed a 5-fold cross-validation experiment by dividing the collection into 5 stratified folds with nonoverlapping BioProjects. This constraint means that a sink is classified without any other samples from the same BioProject in the sources. This data stratification is relevant because it controls for the possible bias that might come from classifying a sink that is similar to the sources simply because they come from the same sequencing initiative and not because there is some underlying biological similarity between the samples (see Figure 12 in Supplementary File for visualisation of the data splitting).

decOM outperforms mSourceTracker and FEAST in each of the five sink/sources folds for performance metrics such as Accuracy, Precision, Recall and F1 Score (see Figure 2) and when metrics are averaged across groups (see Table 2 in Supplementary File). The performance estimates dropped with respect to the leave-one-out MST, which is expected since cross-validation results give a less biased estimate of the model (see also Table 1 and 2 in Supplementary File).

**Table 2:** Performance of decOM in the aOral validation set. As only one class is present in the validation data set (aOral), performance is measured using precision for this highly imbalanced setting.

| Method | Recall |
| --- | --- |
| **decOM** | **0.8654** |
| FEAST | 0.6692 |
| metaSourceTracker | 0.6346 |

#### Validation set

We evaluated decOM in an external validation set with 254 aOral samples that were present in the Ancient-MetagenomeDir [1] but were not part of the matrix of sources previously described. Samples in the validation set belonged to 6 different BioProjects and ranged from 100 to 14800 years old. Furthermore they were isolated from 12 different countries in mostly 2 continents. For more information regarding the metadata of the samples in the validation data set see Supplementary Figures 5 and 6.

Here also decOM outperforms mSourceTracker and FEAST by classifying most of the samples as aOral. See Table 2 for results in the validations set of only aOral samples.

#### Simulated data set

As a final experiment we tested each of the methods on a simulated ancient dental calculus metagenome generated by other authors [33]. A mock oral microbial community is created using representative genomes of microbes found in the human oral microbiome, further processed to appear similar to an ancient metagenomic sample. As in the validation set, we estimated the source environment contribution of the aOral, mOral, skin and sediment/soil microbial communities by using the samples from the 360 collection as sources. Results for all methods are in Figure 3. Given that the synthetic metagenome comes from an uncontaminated mock oral microbial community that has been adapted to appear similar to an ancient calculus sample the expected content is to be 100% oral, decOM provides the highest estimation of oral contribution (ancient or modern), followed by mSourceTracker and lastly by FEAST. We encountered reproducibility problems for FEAST that are further explained in the Supplementary Figure 7.

#### Running times

We measured the running time for decOM and mSource-Tracker using 250 GB of memory and 10 cores. FEAST did not allow for multithreading. We estimated the time it takes to produce an input matrix for each of the methods (whether it is a taxonomy-based clustering table or k-mer matrix of sources). We also estimated the time it takes to analyse a new sample by splitting the process in two steps: the time it takes to produce a new vector to represent the sample, and the time it takes to perform MST. For the two previously mentioned steps, the average running time was estimated on the 254 samples from the validation set. The consolidated running times can be seen in Table3. decOM is an order of magnitude faster than the two other methods for creating a source matrix and producing a new vector. All methods show comparable running times when performing the MST step.

**Table 3:** Running times of MST. Wall-clock time was measured in different parts of the pipeline: Time to build the input matrix, time to produce a new vector from an input FASTQ file and time to perform the MST of one sample. Except for the process named “Build source matrix”, the average time was estimated on the results from the validation set. MST done by FEAST does not allow for multithreading and was run using 2GB of memory and 1 core, whereas mSourceTracker can not split one sink into multiple jobs, so 1 core and 250GB of memory were allocated for each sink. Every other process was run using 250GB of memory and 10 cores. Results for decOM are presented in bold.

| Method | Process | Time (h) |
| --- | --- | --- |
| **decOM** | Build source matrix | **6.60** |
| **decOM** | Produce new vector | **0.04** |
| **decOM** | MST | **0.02** |
| FEAST | Build source matrix | 99 |
| FEAST | Produce new vector | 0.28 |
| FEAST | MST | 0.07 |
| mSourceTracker | Build source matrix | 99 |
| mSourceTracker | Produce new vector | 0.28 |
| mSourceTracker | MST | 0.01 |

#### Ancient oral metagenomic samples come from various environments (multi-source)

After predicting the metadata class of each of the 360 samples in the collection, we also plotted the source proportions according to the estimation done by de-cOM, mSourceTracker and FEAST (Figure 5). The proportion bar plots for mSourceTracker and decOM are visibly more similar to each other than to FEAST, which seems to output more variable results.

**Figure 5:**
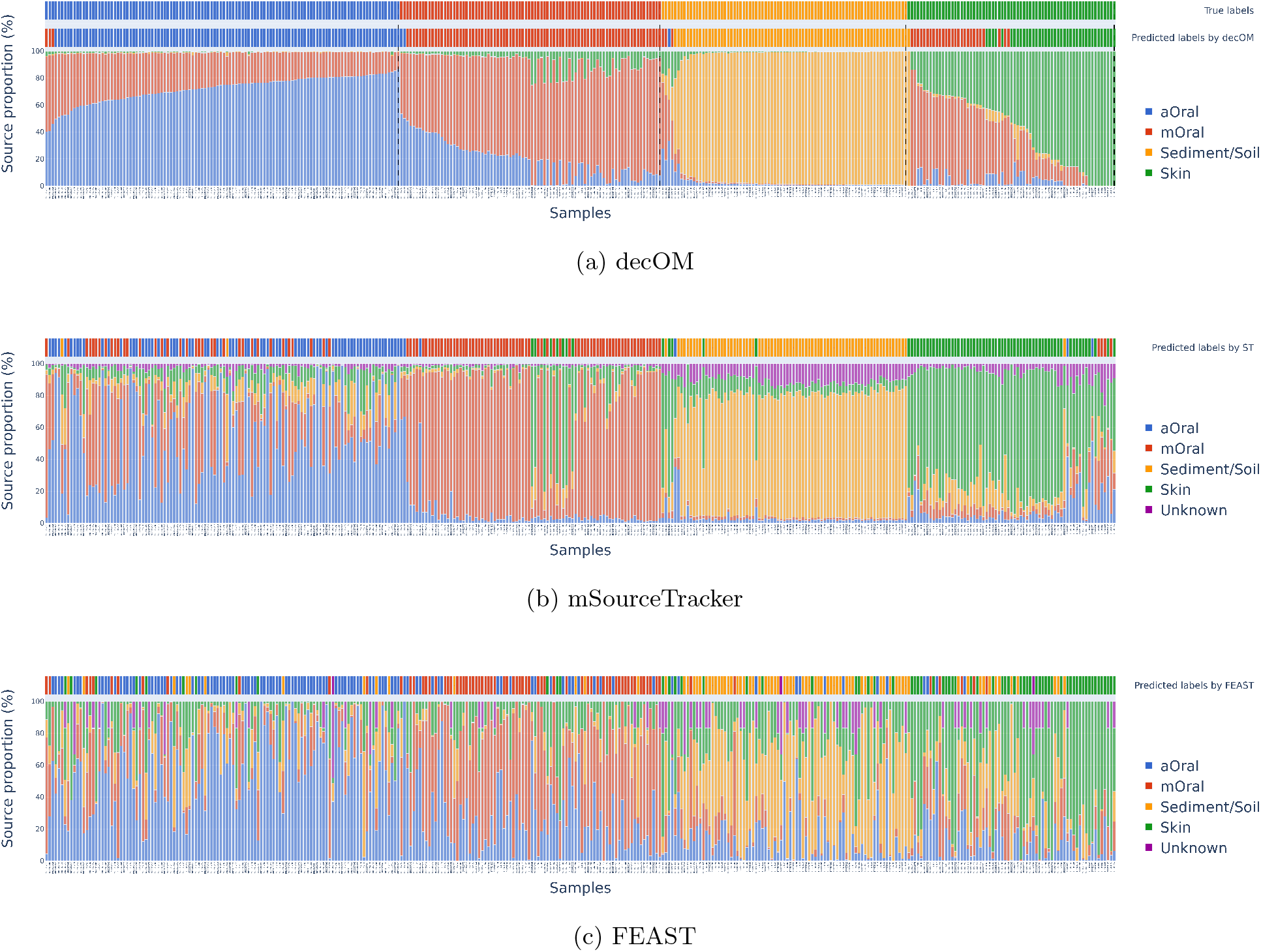
Bar plots of the source environment contribution on each sink after the leave-one-out experiment as estimated by decOM, mSourceTracker and FEAST. Samples in Figure 5a are first sorted by true label, and then sorted by ascending order of the proportion value for such label. Sample order in the x axis for Figures 5b and 5c is sorted according to the order from 5a

According to the estimation done by decOM, there are 4 main predicted groups in the collection with distinct source composition as seen in Figure 5a: there is a group of samples that have a higher sediment/soil content, another class of samples with a higher skin content and with a considerable presence of mOral k-mers, a third group that corresponds to the aOral samples and that also share a part of the mOral content.

Finally, there is a fourth group of samples in which the contribution of the mOral sequences is considerably higher, however these samples also have some k-mers in common with the skin and aOral metagenomic samples.

In additional analyses (see Figure in Supplementary File 15), we divided the samples after decOM’s MST estimation into two categories: samples that come mostly from one source environment (mono-source) or samples that come from several environments (multi-source). In addition to the hard label assigned by decOM, we further categorised the classification of each sample, qualifying the upper quartile (*>* 75%) of each class as mostly mono-source samples, and the first and second quartile (*<* 75%) as samples of diverse origins (more contaminated). According to this threshold, there are 78 mono-source samples (22% of the total collection). These are samples whose recovered label corresponds to the label predicted by decOM, and which are not as contaminated by other sources. A collection of low-contaminated and mono-source samples as this could be used as a high-quality multi-class data set of aOral (36%), mOral (27%), sediment/soil (24%) and skin (13%) for benchmarking with a relatively low imbalance (see Figure 16 in Supplementary File). Interestingly, 91% of the samples we call mono-source are also correctly predicted by mSourceTracker and 78% are correctly predicted by FEAST (Figure 17 in Supplementary File). Nearly a quarter of the aOral samples in the collection have contamination levels that are low enough to have them categorised as mono-source, while the remainder of the ancient oral samples, as expected, have varying levels of contamination.

## Discussion

We have proposed and evaluated decOM as a tool predict the metadata class of a given metagenomic sample by using a Microbial Source Tracking framework, in order to help paleogeneticists better assess the source content of their ancient samples. Because it was built using a Microbial Source Tracking framework, it can also help determine the composition of any other microbial community (not necessarily ancient or of oral origin), which is a common question in microbiome studies. Let us clarify that our goal is not to define an ancient oral microbial community *per se*, but to give the user an indication on the quality of their sample in terms of ancient genetic material. We leave for immediate future work the extensions of decOM to other MST tasks, which could be readily done by creating a k-mer matrix of metagenomic samples of interest with their associated labels and estimating the source proportions using decOM.

The utility of decOM was established on a collection of aOral metagenomic samples and their possible contamination sources, in a leave-one-out set up experiment where every sample was compared against all others. To control for an overly optimistic performance, we performed a stratified 5-fold cross-validation experiment making sure all the samples from the same BioProject belonged to the same fold. Finally, decOM was tested on an external validation data set of 254 aOral samples that were not part of initial collection of metagenomic aOral samples and metagenomes of other contaminants and in a simulated ancient calculus metagenome. We acknowledge that our method would classify the synthetic sample tested on this paper as an mOral sample instead of aOral despite having predicted the largest proportion of aOral source contribution when compared to mSourceTracker or FEAST. However, considering decOM has already proven to be useful on real data, we leave further tuning of the method on synthetic data to be part of the upcoming work. In almost every setting, decOM outperformed two of the most widely-used techniques in the field of MST in the multi-class classification task of predicting the label of a metagenomic samples as the source environment with the highest proportion.

Ideally we would test decOM on a collection of ancient oral samples with known proportions for each source environment, unfortunately, to our knowledge, such a data set does not exist. The task of creating a synthetic data set with such characteristics poses additional challenges regarding how to avoid overlapping species (originating genomes) between each source environment, and would ultimately not be a good representation of a real sample. For this reason we focused on the evaluation of each method by using the meta-data class prediction of a hard label rather than by confirming the proportion predictions were the most accurate.

It could be argued that the lower performance of mSourceTracker and FEAST compared to decOM in the multi-class classification task described in this study was due to limitations of the input taxonomy-based clustering table given to the methods. Better results might be achieved by using a larger database or a tool other than Kaiju to estimate taxonomic abundances. To evaluate this, we conducted an additional experiment in which we constructed another taxonomy-based clustering table with KrakenUniq [34] (see Supplementary File, Section 2). Results in this paper are shown only for the taxonomic abundance profile based on Kaiju, which can also be replicated using public data sets and which, in any case, yielded the best results for the competing methods. The results for the taxonomic abundance profile constructed with KrakenUniq are shown in the Supplementary File information (see Figure 10 and 11).

An important hyperparameter of our model is the size of the input k-mer matrix *M*_*s*_. We explored the effect of using multiple partitions on the performance metrics for the single- and 5-fold cross-validation experiment, but to speed up computations and reduce the memory required, we decided to use only one partition (0.1% of the total k-mer found by kmtricks). Remarkably, the performance of decOM is still better than that of competing methods (see Figures 13 and 14, Table 1 and 2 in the Supplementary File). In the future it would be interesting to study the impact on the classification performance of varying the hyperparameters for the construction of the k-mers matrix, such as the size of the k-mers, minimum recurrence or minimum abundance.

## Conclusions

We propose a novel and reference-free method to perform Microbial Source Tracking and predict the meta-data class of a given (meta)genomic sample. We tested our method on a collection of real metagenomic data sets of aOral origin and its possible contaminants and provided an estimation of the contribution of each source environment on each sample. We anticipate that the incorporation of decOM into paleogenomic analyses will prevent erroneous results and help identify contaminated metagenomic samples and ensure their validity.

## Appendix

## Abbreviations

## Availability of data and materials

The data sets analysed during the current study are available in the repository for decOM [30]. Additional information on the version of the software used, as well as explanation on how to access the run accessions codes for the sources and validation set are present in the Supplementary File.

## Ethics approval and consent to participate

Not applicable.

## Consent for publication

Not applicable.

## Authors’ contributions

## Tables

### Additional Files

Additional file 1 — Supplementary File

Supplementary File (PDF) with supplementary figures and tables cited throughout the text.

