## Supplementary File for "decOM: Similarity-based microbial source tracking of ancient oral samples using k-mer-based methods"

Duitama González, C.

August 12, 2022

**4 Software versions and run accession codes of samples used** **7**

**5 Definition of performance metrics** **8**

### 1 Kaiju taxonomic-based clustering table

Reference database NCBI BLAST nr+euk (2021-02-24 release), is a 64GB non-redundant protein database of bacteria, archaea, viruses, fungi, and microbial eukaryotes can be downloaded at `https://kaiju.binf.ku.dk/database/kaiju_db_nr_euk_2021-02-24.tgz`.

To download and unzip the DB:

```
wget https://kaiju.binf.ku.dk/database/kaiju_db_nr_euk_2021-02-24.tgz
tar -xf kaiju_db_nr_euk_2021-02-24.tgz
```

For every sample analysed (depending wether it single-end or paired-end):

```
kaiju -t nodes.dmp -f kaiju_db_nr_euk.fmi -i reads.fastq [-j reads2.fastq]
kaiju2table -t nodes.dmp -n names.dmp -r genus -o kaiju.table input1.tsv [input2.tsv ...]
```

Then simply parse the results from the resulting .tsv files in an OTU table format suitable for FEAST or SourceTracker.

The results presented in the manuscript were obtained with the taxonomy abundance profile built with this reference database.

#### 2 KrakenUniq taxonomic-based clustering table

The commands used to produce the database using with KrakenUniq were:

```
krakenuniq-download --db DBDIR taxonomy
krakenuniq-download --db DBDIR --dust refseq/bacteria refseq/archaea
krakenuniq-download --db DBDIR
refseq/vertebrate\_mammalian/Chromosome/species\_taxid=9606
krakenuniq-download --db DBDIR refseq/viral/Any viral-neighbors
krakenuniq-download --db DBDIR --dust microbial-nt
krakenuniq-download --db DBDIR contaminants
```

For every sample analysed:

```
krakenuniq --db DBDIR --threads 10
--report-file report_files/{}_unmapped.tax.report.tsv.gz
--output output_files/{}_unmapped.report.tsv.gz --gzip-compressed
--fastq-input {}
```

The symbol {} in this section correspons to a run accession (sample name)

### 3 Commands used to perform Microbial Source Tracking

The file *metagenome_OTU.txt* corresponds to the species abundance profile (taxonomic-based clustering table) that results from parsing the results from Section 1 or 2.

The file *map.txt* is the additional metadata file used as input by SourceTracker and FEAST with three columns (at least): Sample ID with the unique identifier for each sample, SourceSink with the assignment of the sample to either the category source or sink, and finally a column with the name of environment from which each source comes (NA for sink). See documentation of each specific method for their input formats (SourceTracker : `https://github.com/biota/sourcetracker2` , FEAST : `https://github.com/cozygene/FEAST`)

The symbol {} in this section correspons to a run accession (sample name)

#### 3.1 FEAST

The script *Run_FEAST.R* :

```
#!/usr/bin/env Rscript
args = commandArgs(trailingOnly=TRUE)

#Load libraries
library(FEAST)
library(readr)

#Parse arguments
map <- Load_metadata(metadata_path=args[1])
metagenome_OTU <-  Load_CountMatrix(CountMatrix_path=args[2])
accession<-args[3]
dir_path<-args[4]
setwd(dir_path)

FEAST_output <- FEAST(C = metagenome_OTU, metadata = map,
                      different_sources_flag = 0,outfile=accession,
                      dir_path = dir_path)
```

Can be run from the command line in the following manner:

```
Rscript Run_FEAST.R map.txt metagenome_OTU.txt {} ./
```

#### 3.2 mSourceTracker

```
sourcetracker2 -m map.txt -i metagenome_OTU.txt -o output_{}
```

##### 3.3 decOM

For every sample analysed run the following command:

```
decOM -s {} -p_sources decOM_sources/ -k {}.fof -mem 25GB -t 10
```

See details on the format of the key.fof file in `https://github.com/CamilaDuitama/decOM` depending on whether the sample comes from a single-end or paired-end experiment.

### 4 Software versions and run accession codes of samples used

Versions of the software used:

- Kaiju 1.7.3
- KrakenUniq 0.5.8
- decOM 1.0.0
- FEAST 0.1.0
- mSourceTracker 2.0.1-dev

Accession codes are available in `https://github.com/CamilaDuitama/decOM/tree/master/data`, specifically:

- **Collection_accessions.csv:** Accessions for the collection of 360 metagenomic samples that compose the sources of decOM.
- **ValidationSet.csv:** Accessions for the collection of 254 ancient oral metagenomic samples used sinks in validation experiment with an external data set.

### 5 Definition of performance metrics

Consider TP as number of true positives, FN as number of false negatives, $\hat{y}_i$ the predicted value of the i-th sample and $y_i$ the corresponding true value. If we have $n$ samples:

$$\text{Accuracy}(y, \hat{y}) = \frac{1}{n} \sum_{i=0}^{n-1} 1(\hat{y}_i = y_i) \tag{1}$$

$$\text{Precision} = \frac{\text{TP}}{(\text{TP} + \text{FP})} \tag{2}$$

$$\text{Recall} = \frac{\text{TP}}{(\text{TP} + \text{FN})} \tag{3}$$

$$\text{F1-score} = 2 * \frac{(\text{Precision} * \text{Recall})}{(\text{Precision} + \text{Recall})} \tag{4}$$

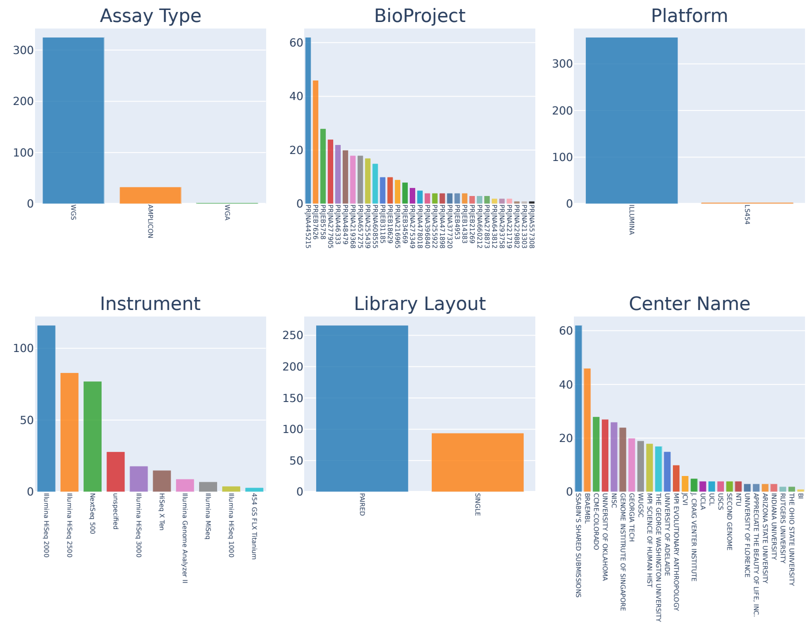

Figure 1: **Metadata barcharts for collection of 360 metagenomic data sets (sources)**. Additional metadata associated to all of the samples in the collection of data sets used to train the method (this includes aOral, mOral, sediment/soil and Skin samples). Barcharts are presented for metadata features such as Assay Type, BioProject, Platform, Instrument, Library Layout and Center Name.

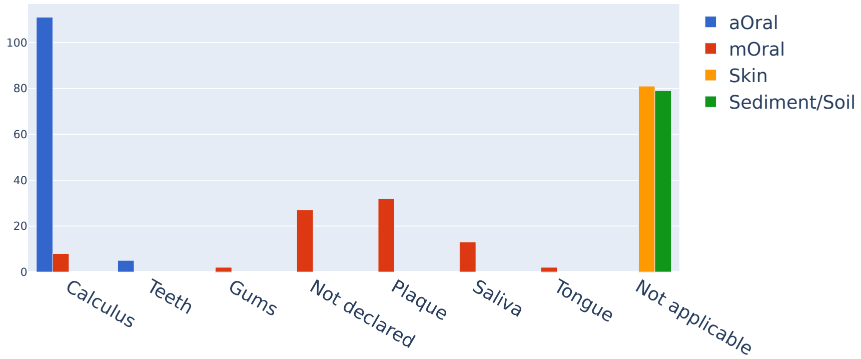

Figure 2: **Isolation source for samples in the collection of 360 metagenomic data sets (sources)**. Barchart colouring corresponds to the original four source environments studied (aOral, mOral, Skin, sediment/soil).

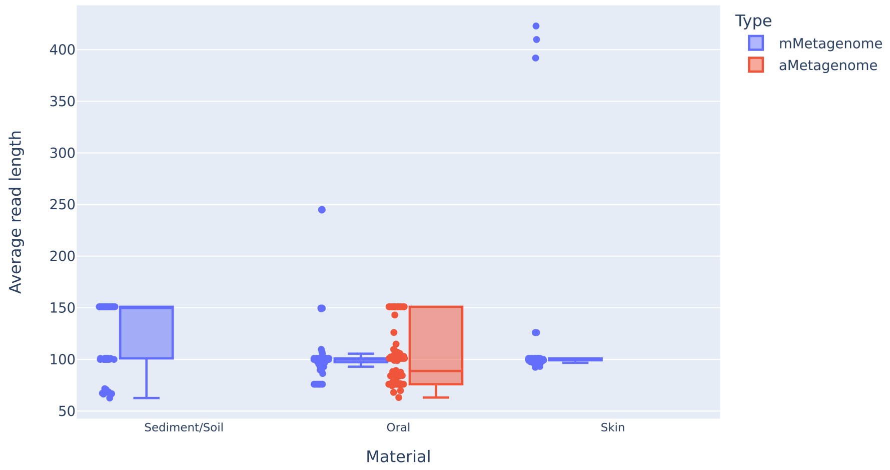

Figure 3: **Average read length for collection of 360 metagenomic data sets (sources).** Samples are grouped by ancient metagenomes (aOral) and modern metagenomes (Skin, sediment/soil, mOral)

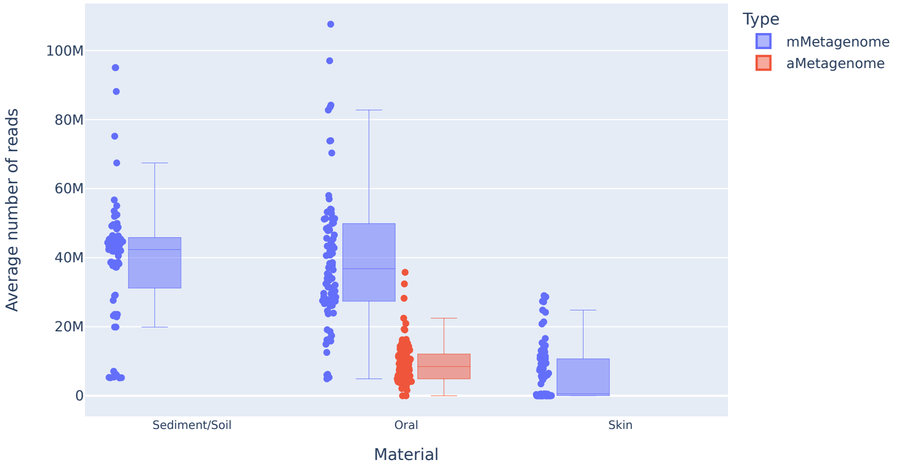

Figure 4: **Average number of reads for collection of 360 metagenomic data sets (sources).** Samles are grouped by ancient (aOral) and modern metagenomes (Skin, sediment/soil, mOral)

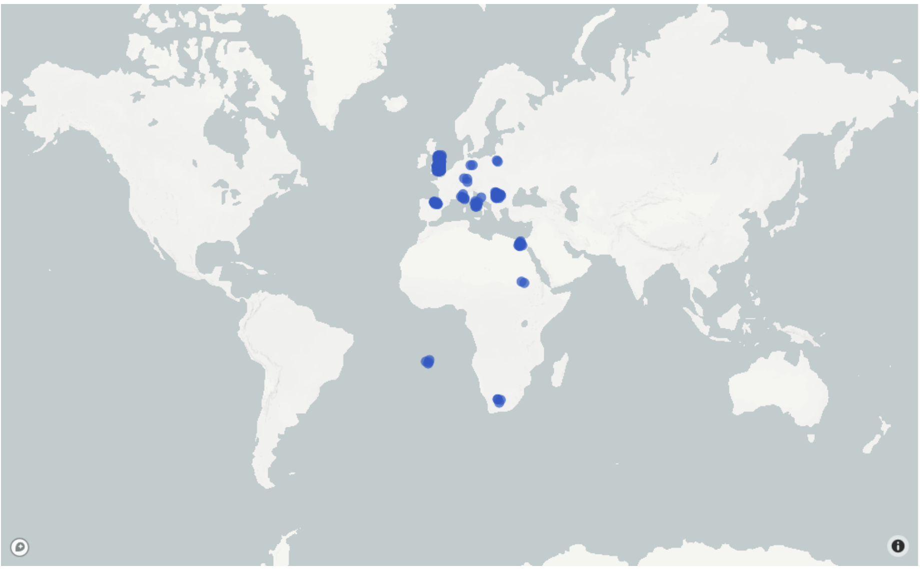

Figure 5: **Origin of samples in validation data set.** All samples in validation set were ancient oral samples obtained from the AncientMetagenomeDir

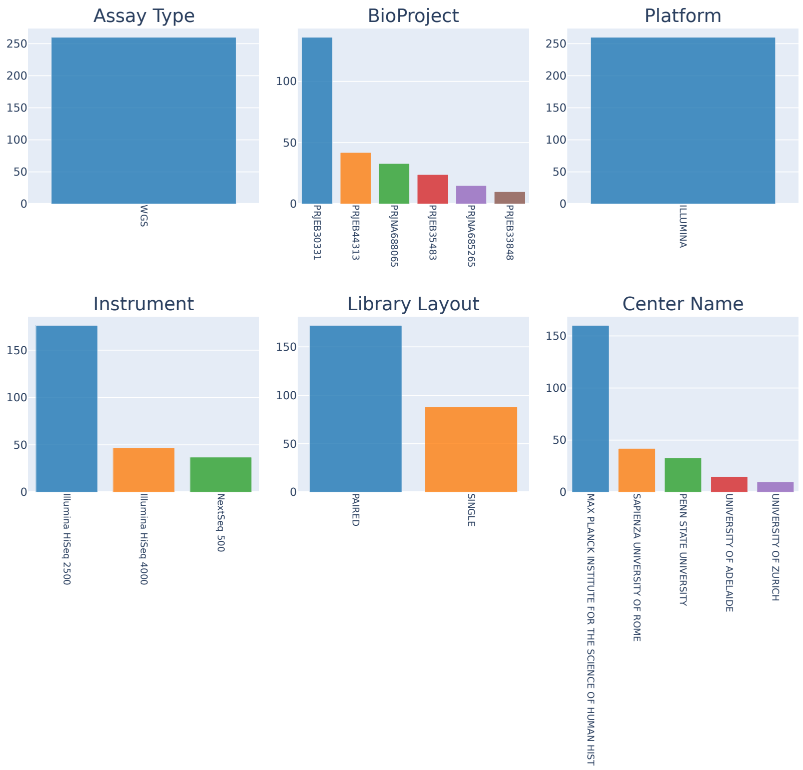

Figure 6: **Metadata barcharts of validation data set**. All samples in validation set are labelled as ancient oral. Barcharts are presented for metadata features such as Assay Type, BioProject, Platform, Instrument, Library Layout and Center Name.

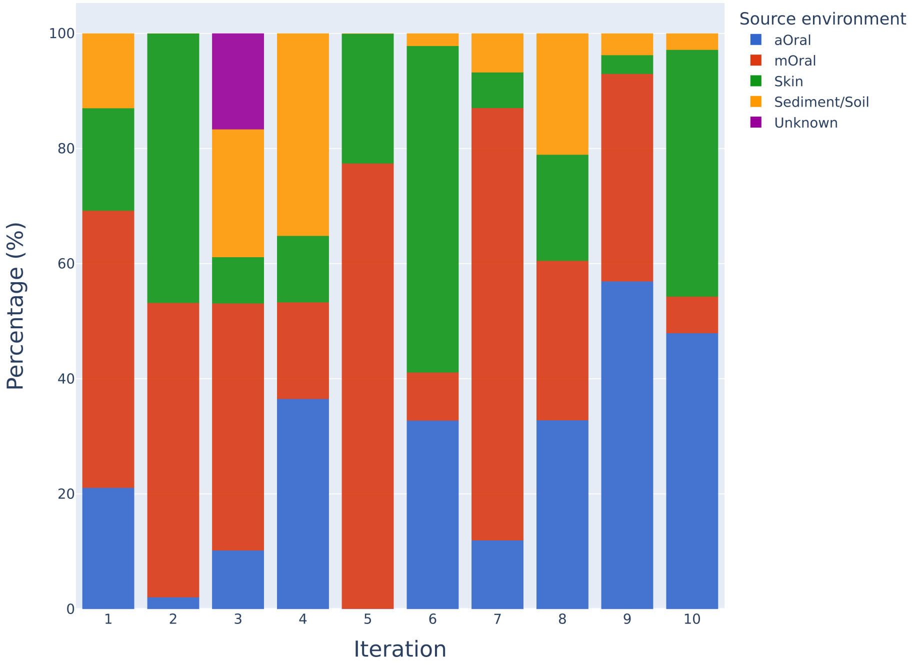

Figure 7: **Microbial source tracking results by using FEAST on the simulated ancient oral data set**. We ran and plotted the source environment bar plots of every output after using FEAST under the same parameters (10 iterations in total) by using as sources all the samples in the 360 metagenomic collection and as sink a simulated ancient calculus data set. As results for this method vary, we selected the first iteration of the outputs to be part of the main text of the paper.

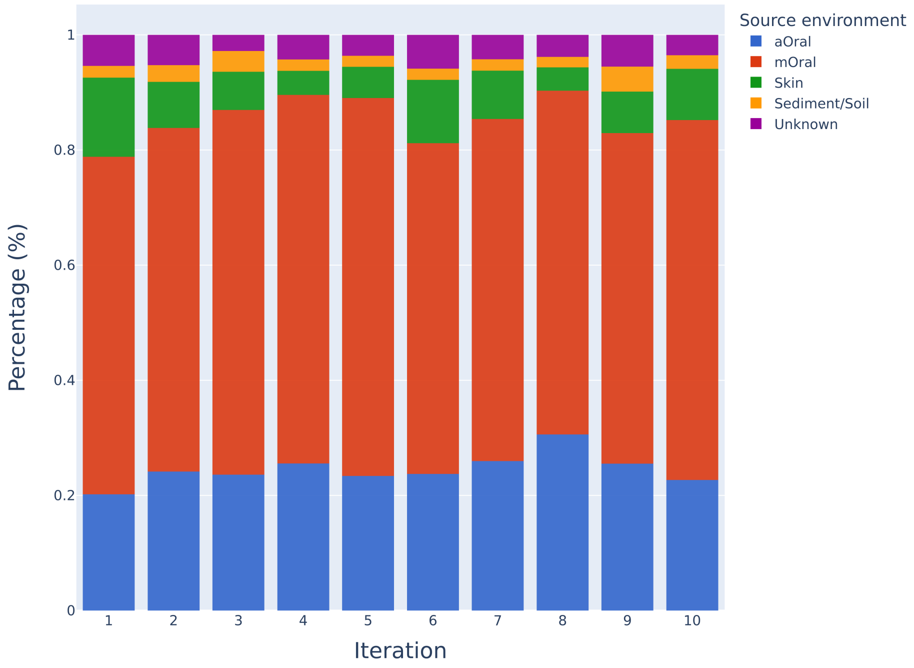

Figure 8: **Microbial source tracking results by using mSourceTracker on the simulated ancient oral data set**. We ran and plotted the source environment bar plots of every output after using mSourceTracker under the same parameters (10 iterations in total) by using as sources all the samples in the 360 metagenomic collection and as sink a simulated ancient calculus data set. See in comparison with Figure 7

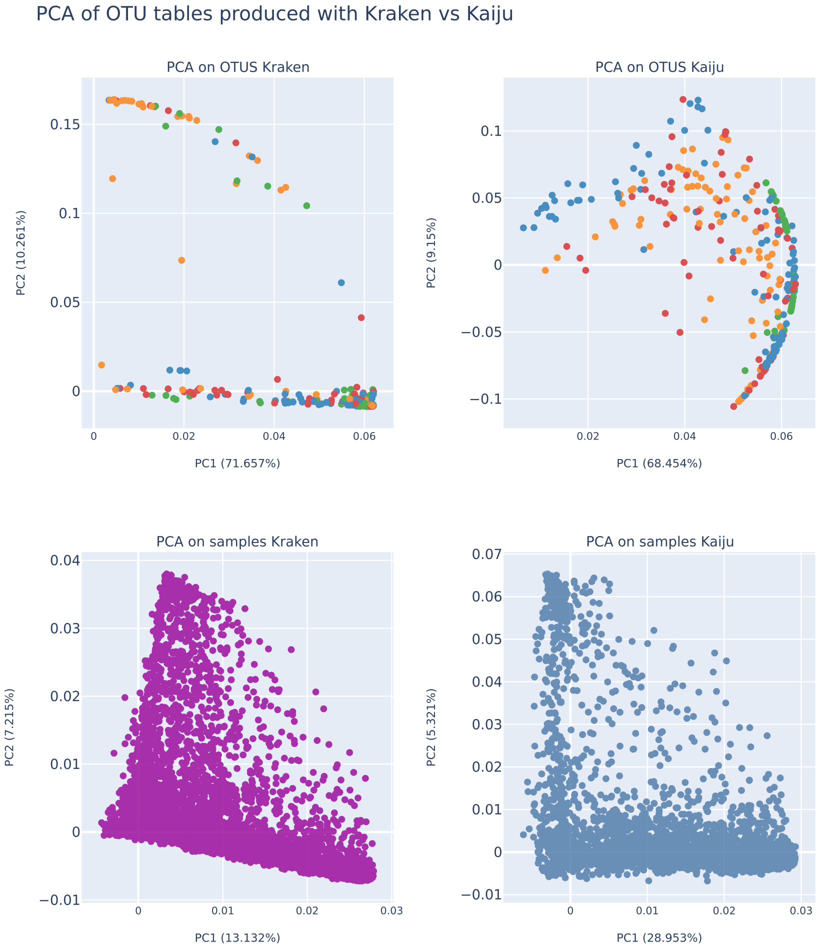

Figure 9: **PCA of OTU table built with Kaiju vs KrakenUniq**. Colour-coding for upper plots: yellow for sediment/soil, green for Skin, blue for aOral and red for mOral samples. Notice that no clear clusters between source environments are seen for either of the OTU tables.

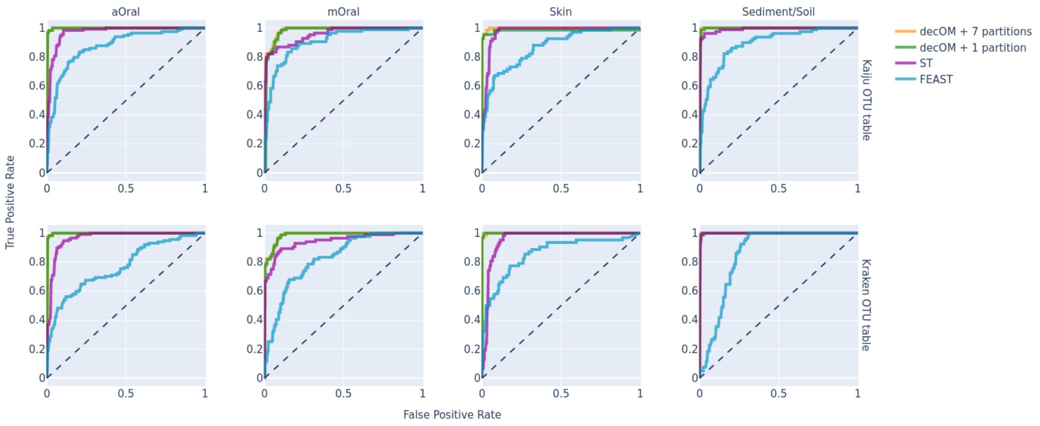

Figure 10: **ROC per class per method.** Each subplot was built using the true and predicted labels for each class, and it includes the curves for every method evaluated. The ROC curve for decOM + 1 partition (yellow) overlaps with the curve for decOM + 7 partitions (green), this is, the performance by using less k-mers is almost not affected.

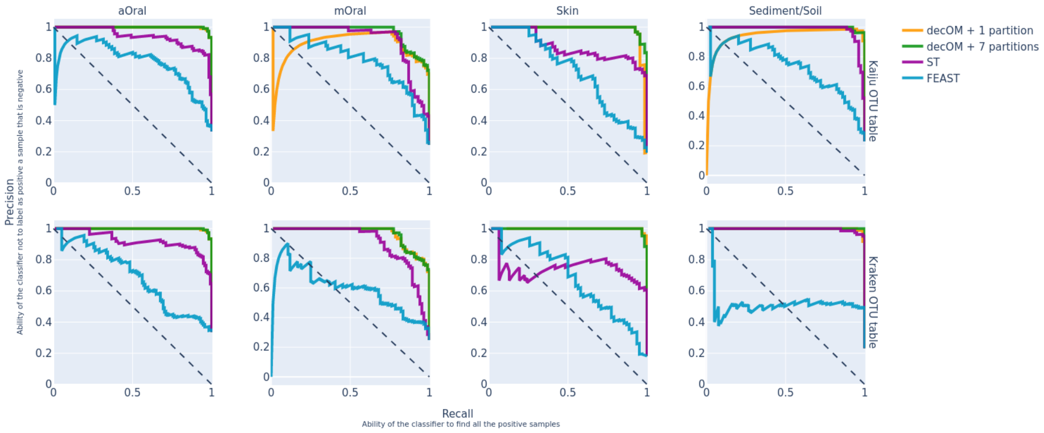

Figure 11: **Precision-Recall curves per class per method.** Each subplot was built using the true and predicted labels for each class, and each subplot includes the curves for every method evaluated. The PR curve for decOM + 1 partition (yellow) overlaps with the curve for decOM + 7 partitions (green), this is, the performance by using less k-mers is almost not affected.

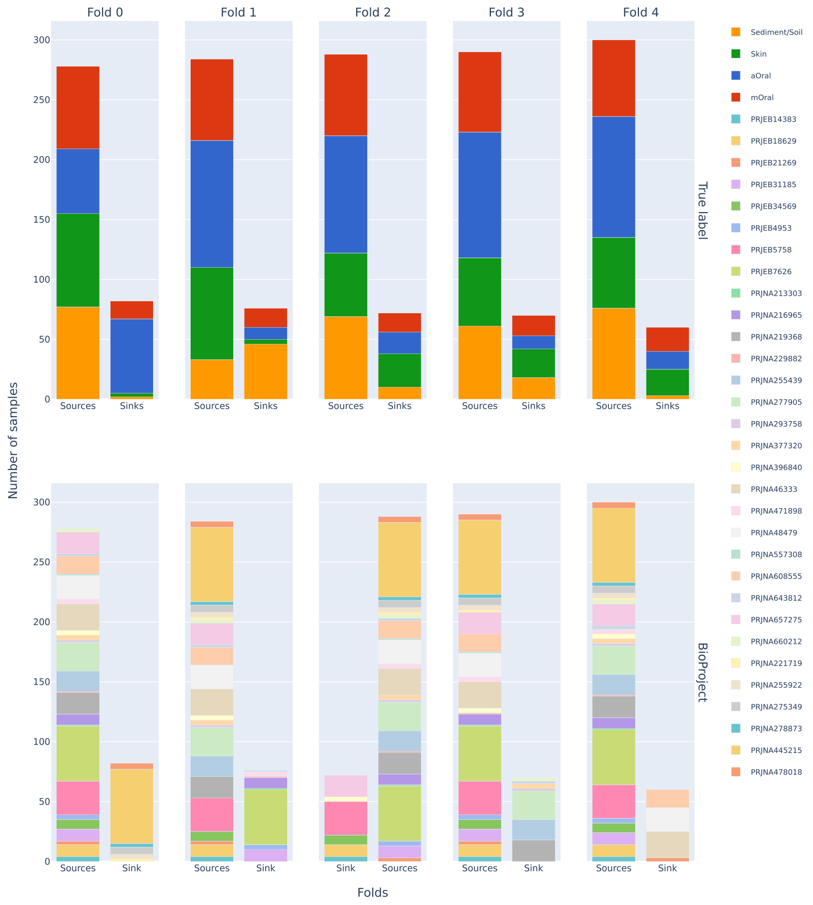

Figure 12: **5-fold cross-validation data split.** Graphic representation of 5-fold cross-validation data split where all samples belonging to the same BioProject are either part of the training or the test set.

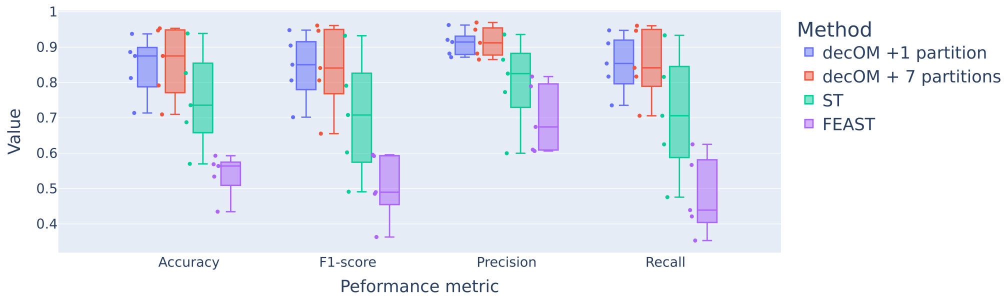

Figure 13: **Performance in 5-fold cross-validation experiment including decOM + 7 partitions.** Box plots for the performance metrics such as Accuracy, F1-Score, Precision and Recall obtained after the the 5-fold cross-validation experiment using decOM + 1 partition, decOM + 7 partitions, mSourceTracker and FEAST.

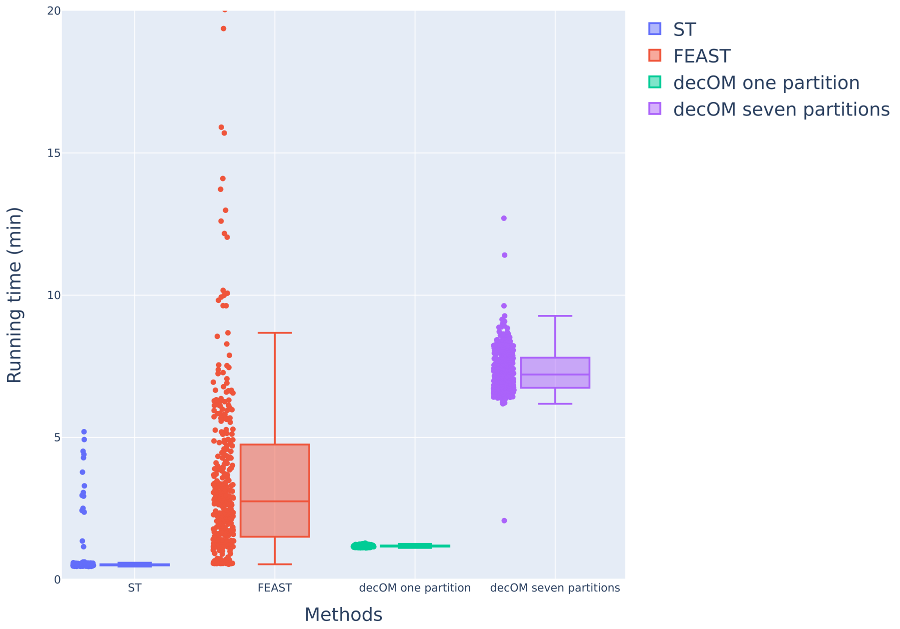

Figure 14: **Running times in leave-one-out experiment.** Box plots built with the running times for each method when analysing each of the samples from the collection (points to the left of the box plot correspond to each measurement). As seen, using one partition is faster on average than using seven partitions.

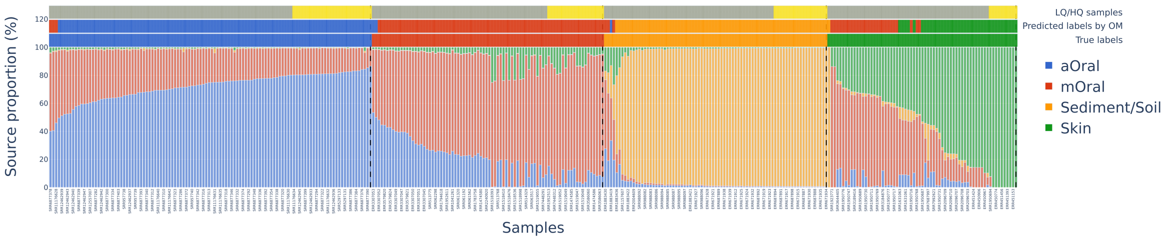

Figure 15: **Estimations of the source contribution to every samples of the 360 metagenomes collection by decOM.** Source estimates considered Skin, Soil, aOral and mOral as possible source environments. The three annotations above the bar chart for each sample from top to bottom correspond to: The legend is light yellow when the sample is considered to be composed of mostly one source for the label assigned (highest 25% of the class, further categorised as mono-source) or grey if it is contaminated and does not come from mostly one source environment (lowest 75% of the class, further categorised as multi-source). The middle annotation corresponds to the predicted label by decOM. Finally, the annotation on the bottom corresponds to the true label for each sample. Samples were first sorted according to their true label, and inside each class they were further sorted with respect to the source proportion estimation for each class label.

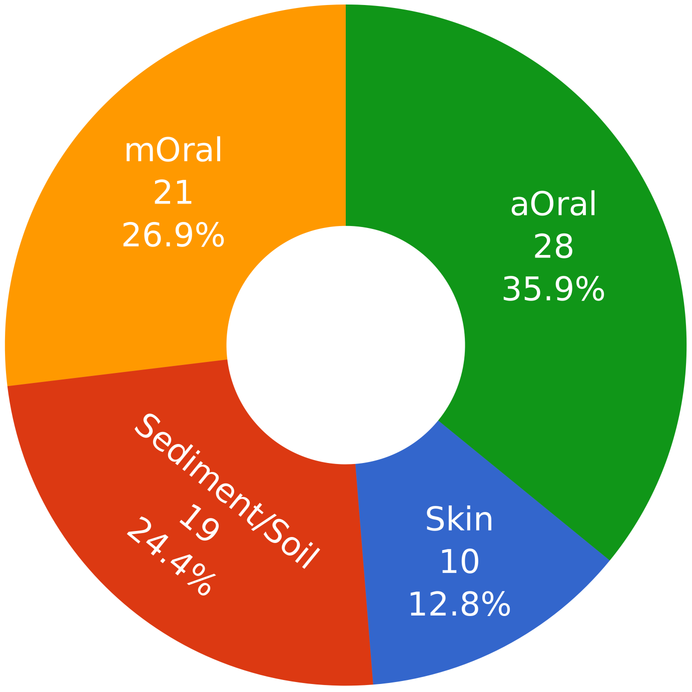

Figure 16: **Class composition of monosource samples as predicted by decOM.** Samples from the collection that we further categorised as monosource samples belong to the classes aOral, mOral, Skin and sediment/soil.

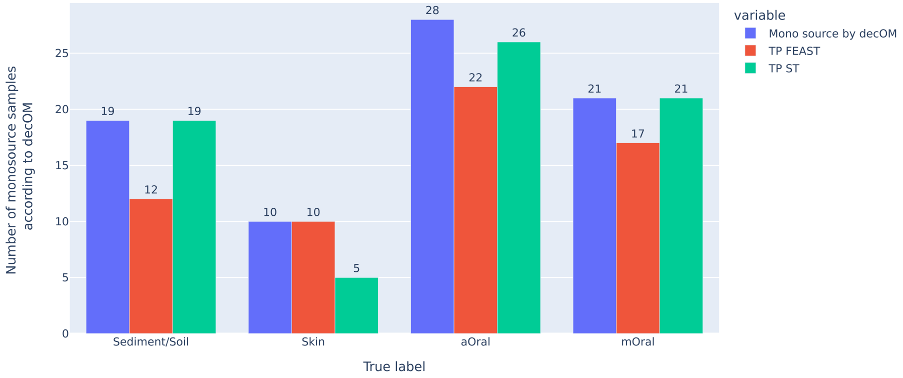

Figure 17: **Percentage of monosource samples according to decOM.** After doing a categorisation of decOM predictions we find some samples in the collection to be composed of mostly one source environment. We distinguish them as monosource. Interestingly, from the monosource samples 61/78(78%) are also correctly predicted by FEAST, whereas 71/78(91%) are correctly predicted by mSourceTracker. TP = True positive

Table 1: **Performance metrics for all three methods compared after leave-one-out experiment.** Performance scores were estimated as the average score across all classes. The result in bold is the one shown in the paper for decOM

| Method | Accuracy | Precision | Recall | F1-score |
|---|---|---|---|---|
| **decOM + 1 partition** | **0.8703** | **0.9184** | **0.8703** | **0.8753** |
| decOM + 7 partitions | 0.8791 | 0.9216 | 0.8791 | 0.8809 |
| FEAST | 0.6816 | 0.5516 | 0.7452 | 0.5479 |
| metaSourceTracker | 0.8388 | 0.8388 | 0.8388 | 0.8289 |

Table 2: **Performance metrics for all three methods compared after stratified 5-fold cross-validation experiment.** Performance scores were estimated as the average score across all classes. The result in bold is the one shown in the paper for decOM

| Method | Accuracy | Precision | Recall | F1-score |
|---|---|---|---|---|
| **decOM + 1 partition** | **0.8450** | **0.9101** | **0.8528** | **0.8421** |
| decOM + 7 partitions | 0.8553 | 0.9156 | 0.8542 | 0.8419 |
| FEAST | 0.5387 | 0.6992 | 0.4809 | 0.5050 |
| metaSourceTracker | 0.7516 | 0.7996 | 0.7111 | 0.7049 |
